# Crosstalk Between Drought-Induced ROS Regulation and Resistance to Xanthomonas oryzae Infection in Rice Plants

**DOI:** 10.1101/2024.10.30.621045

**Authors:** Dhruv Ramu, B. R. Brahmesh Reddy, M. S. Sheshshayee, M. K. Prasanna Kumar

## Abstract

This study investigates the resilience of two genotypes of *Oryza sativa* (rice) with varying levels of tolerance to drought stress, focusing on their production of reactive oxygen species (ROS) and chlorophyll retention when exposed to Xanthomonas oryzae infection. A total of 36 high ATT (AC 39000) and low ATT (BPT5204) plants were subjected to three drought treatments: 100% field capacity (control), gradual drought (reduction to 50% field capacity over 10 days), and rapid drought (immediate reduction to 50%). ROS production was quantified using Evans Blue staining and thiobarbituric acid reactive substances (TBARS) assays, while chlorophyll content was measured using the Arnon method. Membrane damage was higher in the control and rapid stress groups (971 and 1053 ng respectively) compared to gradual drought stress (848.5 ng) on day 12. High-tolerance genotypes demonstrated superior ROS regulation under gradual drought conditions, with the Evans Blue content exceeding that of low-tolerance genotypes by 13 ng. Similarly, chlorophyll retention was significantly higher (p = 0.0005) in high tolerance genotypes (1.166 mg · g^−1^ FW) compared to low tolerance genotypes (0.966 mg · g^−1^ FW). The results indicate that gradual drought stress increases resilience to bacterial infection through enhanced ROS-scavenging mechanisms, which is accentuated in the High ATT genotype, allowing for the development of dual resistant rice varieties capable of withstanding both abiotic and biotic stresses.

## 1 Introduction

Rice, or *Oryza sativa*, is a staple crop for more than half the world’s population and is very important in ensuring global food security [Fukagawa and Ziska, 2019]. However, the challenges posted by abiotic and biotic stresses threaten rice production and yield. Among these, drought-related stress and bacterial infections, particularly by *Xanthamonas oryzae* are particularly problematic. Understanding the crosstalk between abiotic stressors and acquired tolerance to bacteria is crucial for developing resilient crop varieties.

Drought stress is increasingly becoming a pervasive issue due to the changing climate, which exacerbates water scarcity in many rice-growing regions. Bacterial blight, caused by *Xanthamonas oryzae* pv. *oryzae* (Xoo) is known to occur in epidemic proportions around the world [Gnanamanickam et al., 1999].

We expect that the high-tolerance genotype (AC39000) will exhibit more efficient ROS regulation and higher chlorophyll retention compared to the low-tolerance genotype (BPT5204). Plants subjected to gradual drought stress are hypothesized to be more resilient compared to the control group and the rapid stress group due to increased propensity to scavenge ROS.

Reducing the reliance on antibiotics is another significant advantage in building acquired tolerance in rice plants. Excessive use of antibiotics and chemical antimicrobial agents not only poses environmental and health risks but also leads to the evolution of resistant pathogen strains. This has already occurred for certain Xoo strains with resistance towards streptomycin, which is used in the treatment of pulmonary tuberculosis [Xu et al., 2010]. Thus, it is pertinent to study how excessive ROS production can be mitigated in plants to increase their resilience to Xoo.

### 1.1 Stress in Plants

Stress in plants can be formally defined as a “state in which increasing demands on a plant lead to an initial destabilisation of functions, followed by normalisation and improved resistance,” and also, “If the limits of tolerance are exceeded and the adaptive capacity is overworked, the result may be permanent damage or even death.” [Lichtenthaler, 1998]

There are various natural and anthropogenic stressors, but some of the factors of interest are increased temperature, drought, pathogens, and mineral deficiency [Malini et al., 2020, Negrão et al., 2011, Kaur et al., 2021]. The plant stress responses are categorised into the response phase, the restitution phase, the end phase, and the regeneration phase. The response phase observes a decline in vitality and a greater degree of catabolism than anabolism. The restitution phase involves adaptation, repair, and reactivation. Finally, if long-term stress is applied, the end phase involves exhaustion, resulting in possible death [Yavas et al., 2024]. The regeneration phase sees the plant re-establishing normal physiological and metabolic processes.

During photosynthesis, chloroplasts are primary sites for ROS production, especially under conditions that disturb the balance between light energy absorption and its utilisation. Under optimal conditions, the absorbed light energy is used for photochemical conversion in Photosystem II (PSII) and Photosystem I (PSI) [Urry et al., 2020]. However, under stress, excess energy can lead to the formation of ROS. In PSII, singlet oxygen 1 O_2_ is generated when excited chlorophyll molecules transfer energy to molecular oxygen. In PSI, superoxide anions (O_2_^−^) are formed when electrons leak from the electron transport chain to oxygen, which is then rapidly converted to hydrogen peroxide (H_2_O_2_) through dismutation [Foyer and Noctor, 2005].

The specific concentration range of ROS that is conducive to signalling varies depending on the cellular context and the type of ROS. For H_2_O_2_, signalling typically occurs at low micromolar concentrations (1 − 10*µ*M) within cells. In this concentration range, H_2_O_2_ can modify the activity of target proteins, such as transcription factors and kinases.

While ROS serve as signalling molecules, their accumulation beyond certain thresholds (*>* 100*µ*M) can result in the peroxidation of membrane lipids, protein denaturation, and DNA damage [Asada, 2006].

Therefore, plants have evolved an intricate antioxidant defence system that includes both enzymatic and non-enzymatic components to scavenge excessive ROS and maintain cellular homeostasis. Superoxide dismutase (SOD) is the first line of defence against ROS in plants, catalysing the dismutation of superoxide anions. Catalase (CAT) primarily catalyses the decomposition of H_2_O_2_ into H_2_O and O_2_. Other enzymes such as Ascorbate Peroxidase (APX) and Glutathione Reductase (GR) are also involved in this process. In plants, non-enzymatic antioxidants such as carotenoids and tocopherols also prevent membrane damage.

### 1.2 Drought-related Stress

One of the earliest responses to drought stress is the closure of stomata. Under drought conditions, a rapid accumulation of ROS occurs in guard cells, which surround the stomatal aperture. The increase in ROS levels acts as a signal to activate calcium-permeable channels in the plasma membrane of guard cells, leading to an influx of Ca_2_^+^ and Cl^−^ ions into the cytoplasm and an efflux K^+^ ions. This elevation in cytosolic Ca_2_^+^ concentration triggers a cascade of events, including the activation of protein kinases and phosphatases. The resulting osmotic changes cause guard cells to lose turgor pressure, leading to stomatal closure [Xiong et al., 2017]. Furthermore, in some drought-tolerant (High ATT) plant species, the sensitivity of guard cells to ROS is enhanced. Excess ROS can cause lipid peroxidation of thylakoid membranes, impairing the integrity and function of photosystems. In High ATT seedlings, the activity of SOD and APX can increase by up to 50% compared to non-tolerant genotypes [Parida et al., 2022].

### 1.3 Stress from Xanthamonas oryzae

*Xanthomonas oryzae* pv. *oryzae* (Xoo) is a gram-negative bacterium and the causative agent of bacterial blight in rice (Oryza sativa). Xanthomonas oryzae initiates infection by attaching to the rice leaf surface, primarily through hydathodes (water pores) and leaf wounds. The bacteria adhere to the leaf surface using adhesins, polysaccharides, and pili, which facilitate their attachment and help overcome physical barriers on the plant surface [Niño-Liu et al., 2006]. In particular, Xoo utilizes surface-exposed lipopolysaccharides (LPS) and exopolysaccharides (EPS) that contribute to the formation of biofilms on the plant surface. Once adhered, the bacteria penetrate the plant tissue, typically entering the xylem vessels through natural openings. This mode of entry and spread is crucial because the xylem’s function in water and nutrient transport can facilitate the rapid dissemination of the bacteria throughout the plant’s vascular system, resulting in systemic infection.

Xoo has characteristic symptoms of bacterial blight, including leaf yellowing, wilting, and water-soaked lesions that eventually turn necrotic. After entry into the xylem, Xoo establishes itself by producing effector proteins through its Type III secretion system (T3SS), a needle-like apparatus that injects bacterial effectors directly into the plant cells. These effectors, known as Transcription Activator-Like Effectors (TALEs), subvert host cell functions to favour bacterial proliferation. TALEs bind to specific promoter sequences in the plant genome, activating the expression of susceptibility (S) genes. An example is the induction of the OsSWEET family genes, which encode sugar transporters. The upregulation of OsSWEET genes leads to an increased efflux of sugars from the host cells into the apoplast, providing a rich nutrient source that supports bacterial growth and multiplication.

Upon Xoo infection, ROS are primarily generated at the plasma membrane by NADPH oxidases and in organelles such as chloroplasts, mitochondria, and peroxisomes. The H_2_O_2_ produced during the oxidative burst can diffuse through membranes and participate in reinforcing cell walls via the cross-linking of lignin and other structural polymers, thereby restricting pathogen movement. The local concentration of ROS in infected tissues can exceed 10 µM, a level sufficient to trigger programmed cell death (PCD) in neighbouring cells, a process known as the hypersensitive response (HR). The HR is a defence strategy that creates a physical barrier around the site of infection, limiting the spread of the pathogen. Simultaneously, ROS serve as secondary messengers that activate other defence responses, including the induction of pathogenesis-related (PR) proteins, the production of secondary metabolites (e.g., phytoalexins), and the expression of defence-related genes [Ma et al., 2018].

In Oryza sativa, the accumulation of ROS leads to the activation of the WRKY transcription factors, which modulate the expression of defence genes involved in the synthesis of PR proteins and callose deposition. Callose, a *β*-1,3-glucan, is deposited at the site of infection to reinforce the cell wall and prevent further pathogen ingress.

During Xoo infection, the pathogen’s effectors can hijack defence pathways, leading to an imbalance in ROS generation and scavenging processes. For example, the suppression of salicylic acid-mediated defences can impair the activation of key antioxidant enzymes such as superoxide dismutase (SOD) and catalase (CAT), resulting in an accumulation of ROS beyond normal levels [Jiang et al., 2013]. As ROS levels continue to rise, the pool of non-enzymatic antioxidants, such as glutathione and ascorbate, becomes depleted. This is a positive feedback mechanism as it will result in greater ROS production.

## 2 Method

### 2.1 Experimental Design and Procedure

#### 2.1.1 Preparation of Plants

Soil was collected and placed in a hot air oven at 45°C for 48 hours until it was ‘bone dry’. The field capacity (FC) is calculated using formula 1. Subsequently, the soil was evenly distributed in 36 plastic cups, each containing approximately 190 grams (dry weight) of soil. The prewashed rice seeds, belonging to two distinct genotypes (18 replicates each), exhibiting high or low acquired tolerance traits (ATT) to abiotic stress, were planted in these cups, with three seeds per cup. Each genotype group was further divided into three treatment subgroups, representing varying drought stress conditions: Control (100% FC), Gradual Drought Stress, and Rapid Drought Stress, with six replicates per treatment. All cups were initially watered uniformly to maintain 100% FC to promote seed germination.

#### 2.1.2 Drought Stress Induction

After true leaf formation, drought stress was then introduced in the following manner: the control group continued to receive sufficient water to maintain 100% FC throughout the experiment. For the Gradual Stress group, the field capacity was incrementally reduced by 5% each day over 10 days until it reached 50% FC. In contrast, the Rapid Stress group experienced an immediate reduction to 50% FC, maintained by periodic weighing and adding water. These treatments were maintained for a total of 10 days, after which all plants were allowed to recover under optimal watering conditions, restoring FC to 100% for another 10 days.

Drought stress can be induced by varying the field capacity of the plant. The below formula has been utilized:

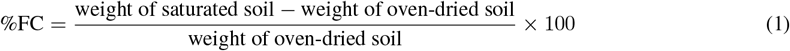

#### 2.1.3 Bacteria Induced Stress

To simulate biotic stress, *Xanthomonas oryzae* was inoculated with the plant. The bacterial stock was stored at -80°C and revived on kanamycin-amended agar medium using the spread plate method. After a 5-day incubation at 32^dss^C, a colony was transferred to nutrient broth and cultured at 32^°^C for 2 days with shaking (150 RPM) until it reached an optical density between 0.6 and 1.0 at 600 nm. A uniform wound was then introduced on one leaf of each seedling by dipping sterilised scissors into the bacterial suspension and making a cut of 2 cm. The plants were kept in an airtight greenhouse at 28^°^C.

#### 2.1.4 Quantification of Reactive Oxygen Species

To assess the physiological responses of the plants, several measurements were conducted. Evans blue staining is used for assessing cell membrane integrity, providing a measure of cell death and oxidative stress using spectrophotometry. It is a negatively charged dye that cannot permeate intact cell membranes. However, during stress conditions, ROS-induced lipid peroxidation can damage the membrane, increasing its permeability. As a result, Evans blue selectively stains dead or damaged cells [Vijayaraghavareddy et al., 2017].

ROS production was evaluated using Evans Blue Staining to quantify membrane damage with 0.25g Evans Blue solution prepared in CaCl_2_ (0.1 M, pH 5.6) and incubated for 1 hour [Yao et al., 2018]. The samples were then thoroughly rinsed with water, after which they were ground in a crucible, treated with 1% SDS, and centrifuged at 12000 RPM for 15 minutes. Absorbance was recorded at 600 nm [Nv et al., 2017].

The Thiobarbituric Acid Reactive Substances (TBARS) assay is utilized for quantifying lipid peroxidation in rice plants by measuring malondialdehyde (MDA), a key breakdown product of polyunsaturated fatty acids [Aguilar Diaz De Leon and Borges, 2020]. A 2 cm leaf sample was taken from each plant and promptly stored in an ice bath, after which it was stored at − 80^°^C. Malondialdehyde (MDA), the by-product of lipid peroxidation, was extracted using 5% trichloroacetic acid (TCA) and centrifuged at 12000 RPM for 15 minutes. To 1 mL of the extract, 2 ml of 0.5% TBA in 20% TCA was added and incubated at 95^°^C for 30 minutes. Absorbance was recorded at 532 and 600 nm [Aguilar Diaz De Leon and Borges, 2020].

#### 2.1.5 Quantification of Chlorophyll Content

Chlorophyll content is an indicator of plant health and photosynthetic efficiency. DMSO acts as an organic solvent, extracting chlorophyll from leaf tissue without altering its structure. The absorbance of the resulting chlorophyll extract is then measured spectrophotometrically and chlorophyll a and b per mg is calculated using the Arnon method formula[Arnon, 1949].

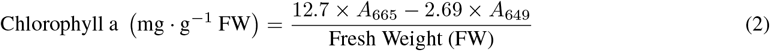

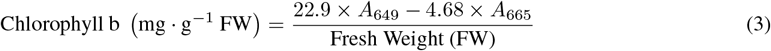

The chlorophyll content was measured by submerging 2 cm leaf samples in DMSO for 24 hours. Then, absorbance of the sample was recorded at 645 and 663 nm and used to calculate the total chlorophyll content using equations 2 and 3.

## 3 Results

### Nomenclature

**Table 1:** Nomenclature of Samples: ATT, Drought Stress, and Bacterial Exposure.

| Sample | ATT Genotype |  | Drought Stress |  |  | Bacterial Exposure |  |
| --- | --- | --- | --- | --- | --- | --- | --- |
|  | High | Low | Control | Gradual | Rapid | Yes | No |
| HCC | ✓ |  | ✓ |  |  |  | ✓ |
| HCF | ✓ |  | ✓ |  |  | ✓ |  |
| HGC | ✓ |  |  | ✓ |  |  | ✓ |
| HGF | ✓ |  |  | ✓ |  | ✓ |  |
| HRC | ✓ |  |  |  | ✓ |  | ✓ |
| HRF | ✓ |  |  |  | ✓ | ✓ |  |
| LCC |  | ✓ | ✓ |  |  |  | ✓ |
| LCF |  | ✓ | ✓ |  |  | ✓ |  |
| LGC |  | ✓ |  | ✓ |  |  | ✓ |
| LGF |  | ✓ |  | ✓ |  | ✓ |  |
| LRC |  | ✓ |  |  | ✓ |  | ✓ |
| LRF |  | ✓ |  |  | ✓ | ✓ |  |

### 3.1 Shoot Length

Across all conditions, high ATT genotypes (average shoot length = 52.6 cm) maintained a longer average shoot length than low ATT genotypes (average = 46.8 cm). The heightened shoot length for high ATT genotypes suggests stronger growth stability under drought conditions. In the gradual drought condition, high ATT genotypes averaged 53.8 cm in shoot length, while low ATT genotypes averaged 48.1 cm. In rapid drought conditions, high ATT genotypes experienced a slight reduction (to 50.9 cm), whereas low ATT genotypes were more affected, dropping to an average of 45.5 cm. This trend indicates that high ATT genotypes better adapt to stress when it is gradual, while low ATT genotypes are more vulnerable to sudden stress changes.

### 3.2 Qualitative Observations

### 3.3 Effect of Bacteria on Chlorophyll Content

As seen in Figure 2, the Low ATT plants were more adversely affected in terms of the formation of lesions. This has also been reflected in the total chlorophyll content. High ATT genotypes began with a higher average chlorophyll content (1.667 mg · g^−1^ FW) compared to low ATT genotypes (1.276 mg · g^−1^ FW) due to known improved resilience against drought stress in the High ATT genotype. By Day 5, chlorophyll content in high ATT genotypes under gradual drought dropped only slightly to an average of 1.569 mg · g^−1^ FW. Under rapid drought, this decreased more sharply to 1.307 mg · g^−1^ FW. Low ATT genotypes experienced a more substantial decrease by Day 5, especially under rapid drought, where levels dropped from 1.276 mg · g^−1^ FW to 1.105 mg · g^−1^ FW. By Day 12, the average chlorophyll in low ATT genotypes under rapid drought dropped further to 0.966 mg · g^−1^ FW, while high ATT genotypes maintained a higher content of 1.212 mg · g^−1^ FW even under similar stress. These differences suggest that high ATT genotypes better preserve chlorophyll under drought and bacterial stress, maintaining photosynthetic activity longer than low ATT genotypes. This trend, particularly in gradual drought conditions, highlights the high ATT genotype’s superior ability to modulate stress responses over time, invariant of whether or not it is exposed to drought or bacterial stress.

**Figure 1:**
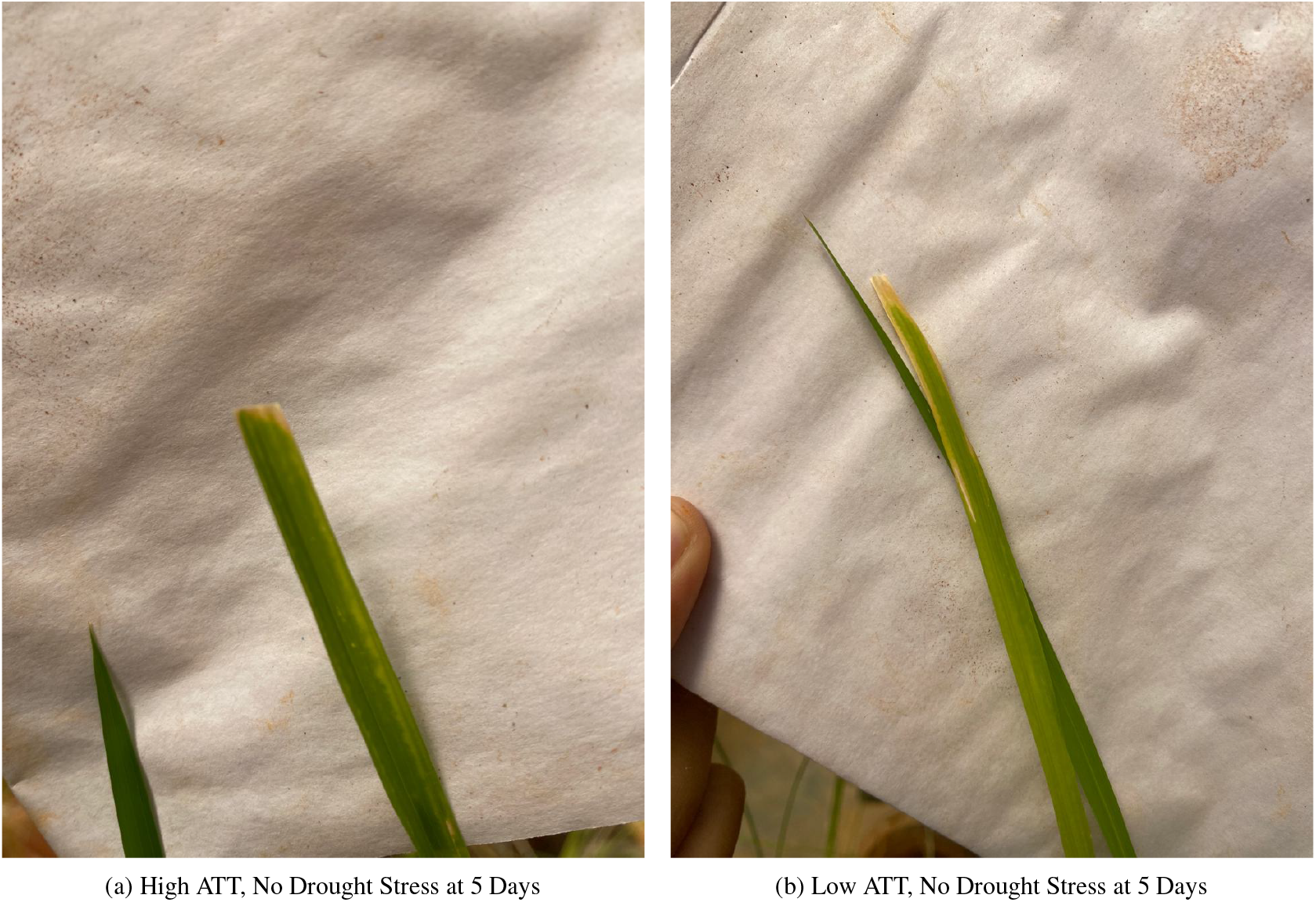
Qualitative comparison of biotic stress on the leaf.

**Figure 2:**
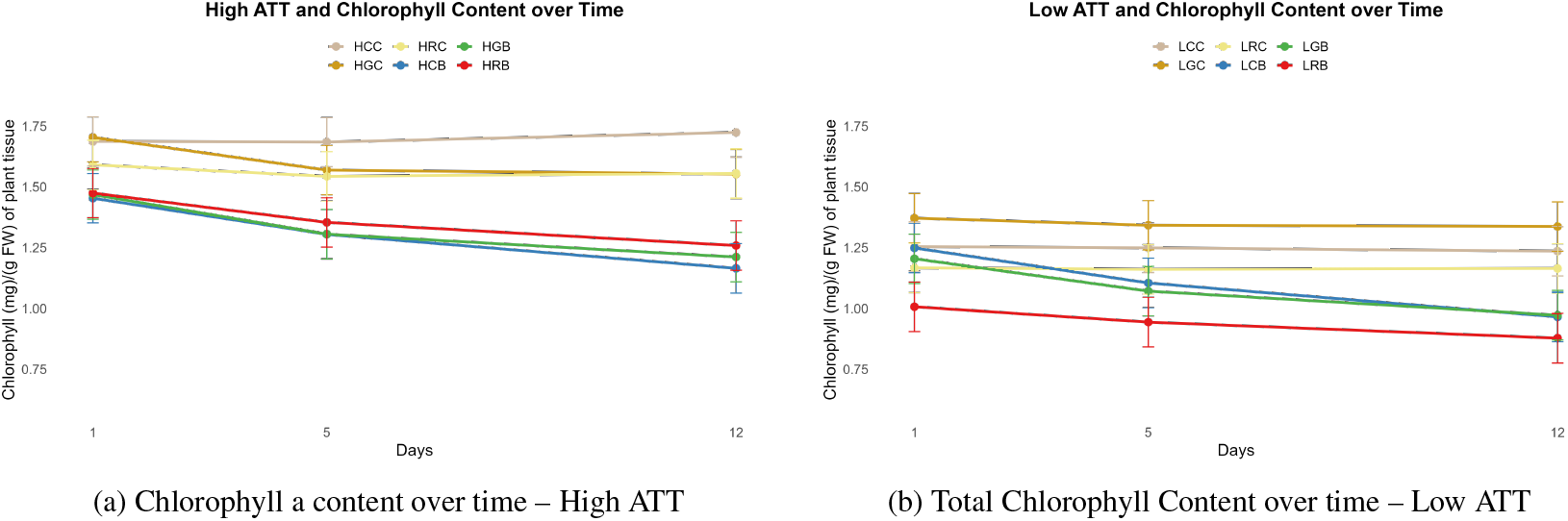
Chlorophyll content over time for High and Low ATT

### 3.4 Effect of Bacteria on ROS Production

High ATT genotypes under gradual drought exhibited consistently lower ROS production across Days 1, 5, and 12, with averages of 268.0, 273.0, and 266.0 ng of Evans Blue, respectively. These stable values indicate minimal ROS-induced membrane damage. Under rapid drought, however, these values rose to 336.5, 751.5, and 848.5 ng by Day 12, highlighting increased ROS accumulation when drought stress was abrupt. Low ATT genotypes, especially under rapid drought, exhibited higher average values starting from 349.0 ng on Day 1 and reaching 975.0 ng by Day 12. However, we show that if drought stress is introduced gradually as seen in Figure **??**, there is comparatively less lipid peroxidation as the MDA content was least in the gradually stressed low ATT genotype (LGB = 0.088 nmol/mg MDA on day 12) compared to the rapid stress and control group.

## 4 Statistical Analysis

The *r* values in Figure 6 were calculated to determine the strength of the correlation of the chlorophyll content over time for the infected treatment groups. It is strongly negative in all of them. It can be seen, however, that it was marginally lower in the gradually stressed group (HGB) compared to the control (HCB).

## 5 Discussion

The results confirmed that the High ATT (AC 39000) genotype demonstrated superior resilience to *X. oryzae* under drought stress compared to the Low ATT (BPT5204) genotype. Specifically, plants in the High ATT group exhibited more effective ROS management, as evidenced by lower levels of ROS accumulation, which was quantified using Evans Blue staining and lipid peroxidation assays (0.791 A in gradual vs 0.914 A in control).

Various studies demonstrate increased susceptibility to pathogens with simultaneous abiotic stress such as high temperature stress in tobacco to Tobacco mosaic virus and in Arabidopsis to *Pseudomonas syringae* [Király et al., 2008, Wang et al., 2009]. It has also been observed that previously drought stressed Phaseolus vulgaris plants were more adversely affected with a decreased transpiration rate, water potential, and relative water content [Mayek-PÉrez et al., 2002].

The acclimation of gradually stressed plants (represented by the greatest increase in membrane damage as seen in Figures 3a and 3b reinforces that abiotic and biotic stress is not independent (paired t-test reveals p = 0.0005). Under moderate drought stress conditions (e.g., 50% field capacity), rice plants were observed to activate a protective mechanism involving the scavenging of ROS through antioxidant enzymes such as SOD and CAT, priming plants for more severe stress conditions [Ramu et al., 2020]. The crosstalk between abiotic and biotic stress responses in rice is also regulated by ABA, SA, and JA.

**Figure 3:**
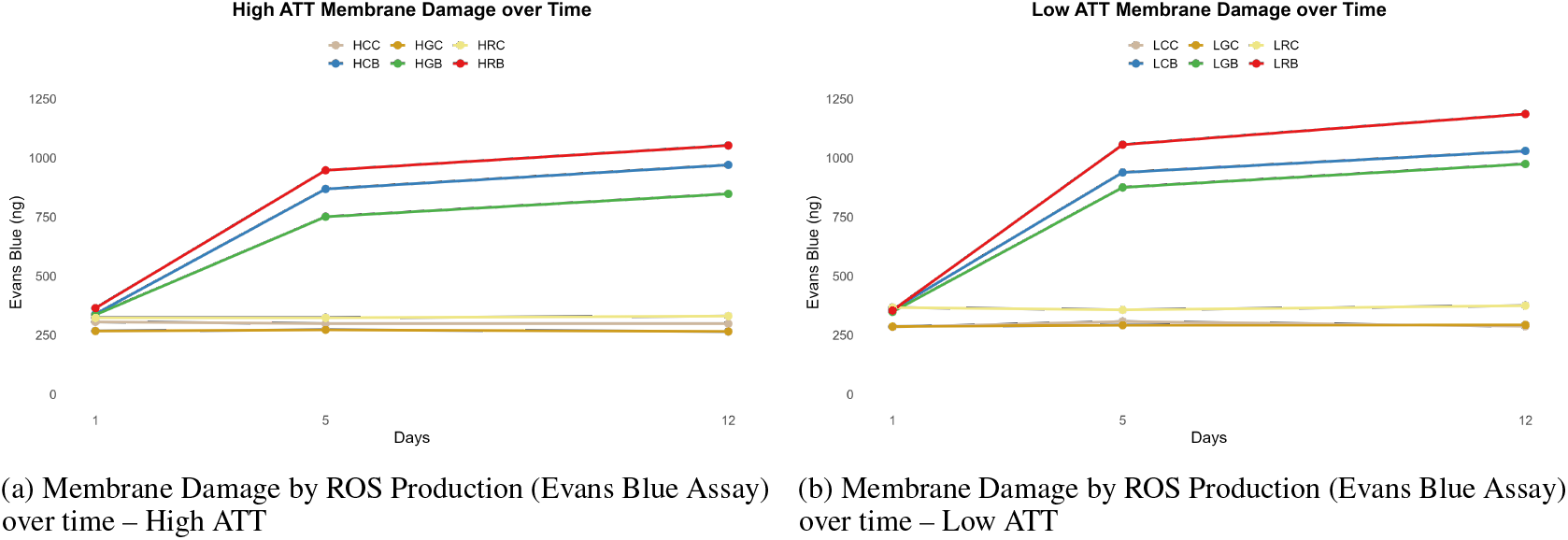
Membrane Damage by ROS Production (Evans Blue Assay) for High and Low ATT over time

**Figure 4:**
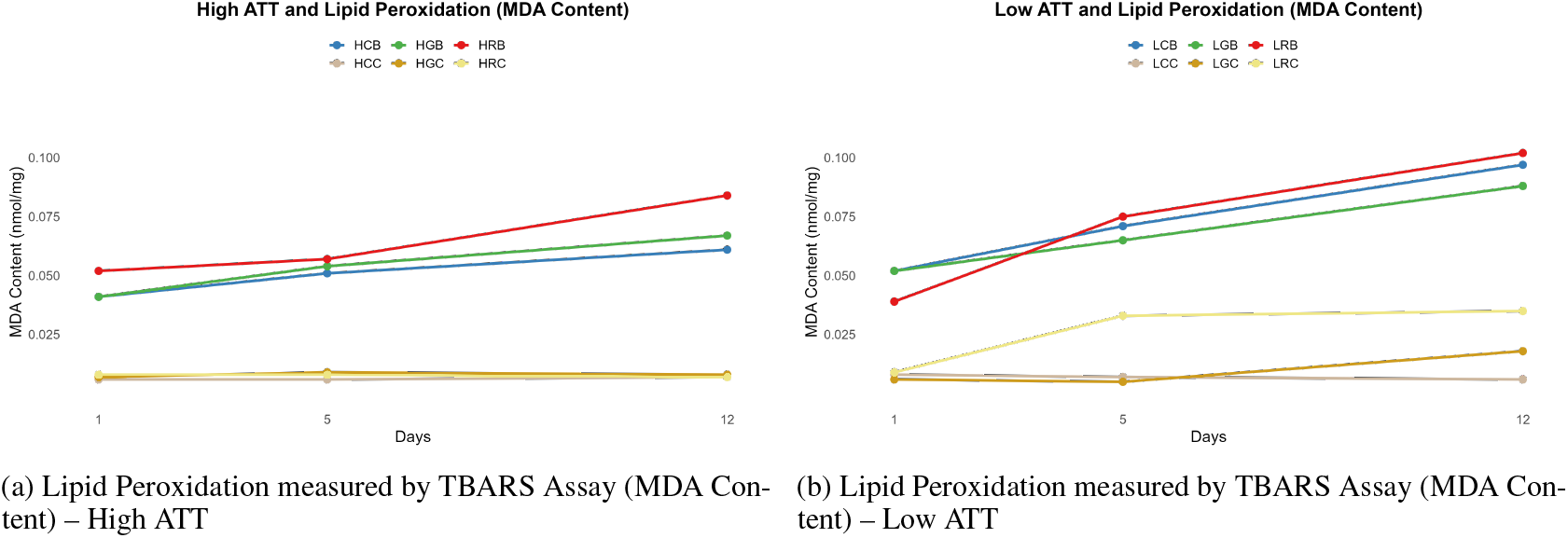
Lipid Peroxidation measured by TBARS Assay (MDA Content) for High and Low ATT

**Figure 5:**
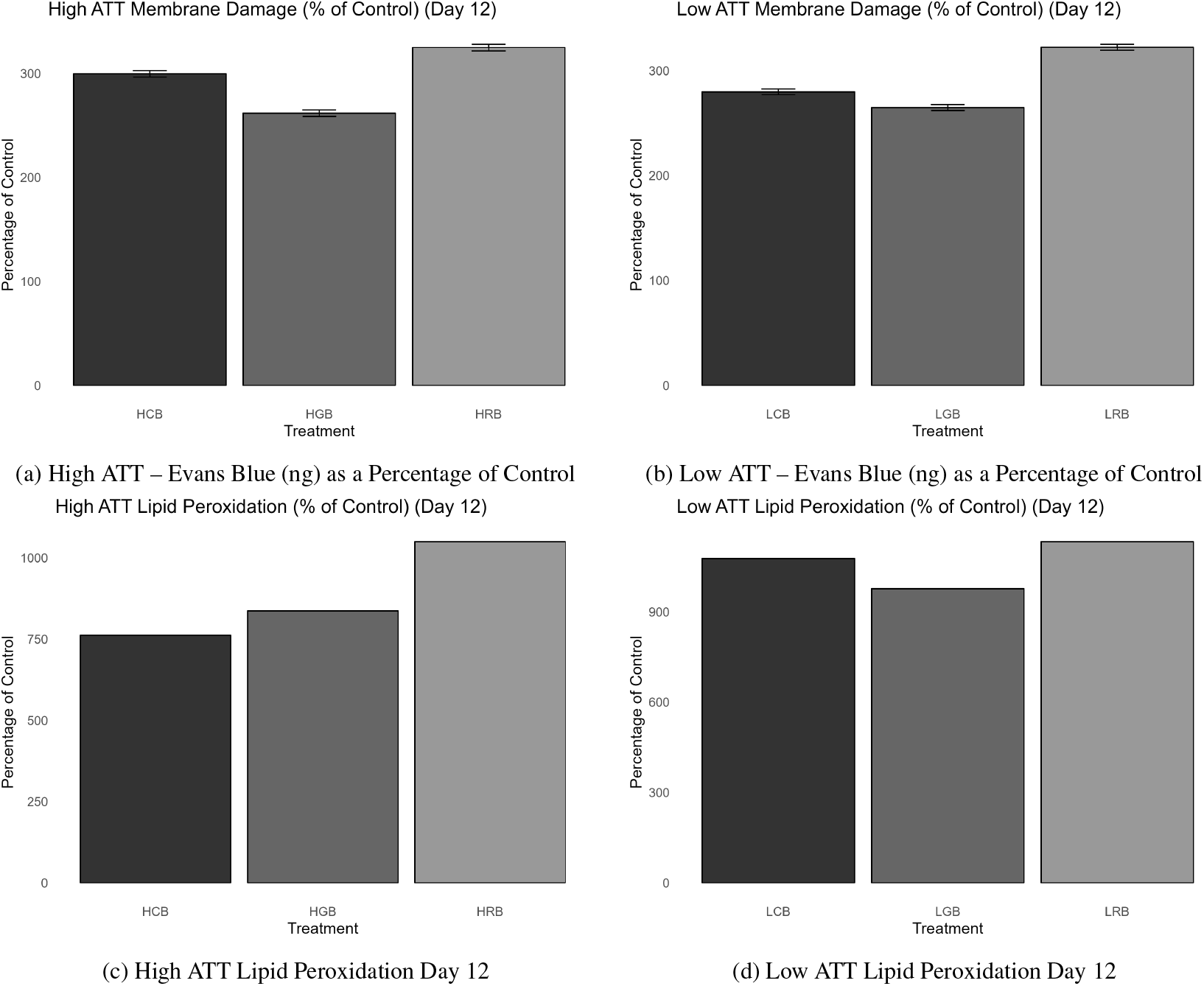
Quantification of ROS with Evans Blue (ng) and Lipid Peroxidation measurements for infected groups against the control.

**Figure 6:**
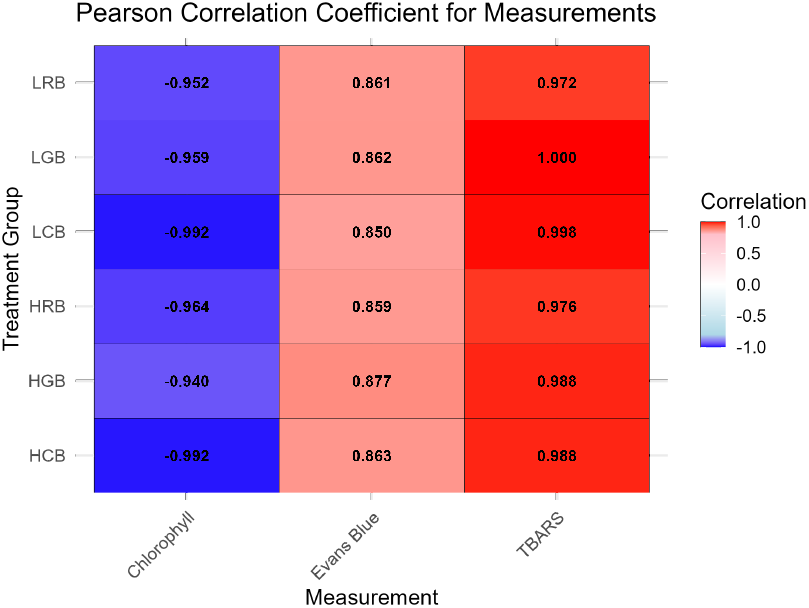
Pearson Correlation Coefficient (*r* value) for the relationship between treatment groups and the dependent variable over time.

ABA is predominantly associated with the plant’s response to drought stress, where it promotes stomatal closure to reduce water loss and induces the expression of drought-responsive genes [Asselbergh et al., 2008]. However, under rapid stress, ABA can reduce the plant’s ability to mount an effective defence response by suppressing the SA and JA pathways that are essential for activating defence-related genes [Ramegowda et al., 2013, Anderson et al., 2004]. Rapid drought stress disrupts cellular homeostasis before adaptive responses can be fully activated, leading to protein denaturation, membrane lipid peroxidation, and chlorophyll degradation, which have been observed in this experiment. There is also a genetic basis for this crosstalk that has been extensively observed in Arabidopsis [Seki et al., 2002, Swindell, 2006, De Vos et al., 2005]. Rice plants pre-exposed to moderate drought stress showed increased upregulation of ascorbate peroxidase, FeSOD, CAT, and aldo-keto reductase at 60% FC which is likely in this experiment as well [Vijayaraghavareddy et al., 2022]. There is also over-expression of the transcription factor OsDREB2A during drought stress, which increases tolerance, but also regulates the expression of genes involved in defence against pathogens, indicating convergence of abiotic and biotic stress signaling pathways [Matsukura et al., 2010, Zhang et al., 2013, M et al., 2011].

The High ATT genotype (AC 39000) has been shown through the TBARS assay to be superior in drought stress management[Lekshmy, 2022], and it appears that this has been conserved for biotic stress as well as seen in Figures 3a and 3b, where the drought stress-free high tolerance group (HCB) observed 0.011 less nmol/mg of MDA. In contrast, the Low ATT genotypes were more susceptible to the combined effects of drought and bacterial infection. These plants exhibited higher ROS accumulation, which contributed to significant cell damage and increased membrane permeability.

## 6 Conclusion

These results have important implications for breeding programmes aimed at enhancing rice resilience to both abiotic and biotic stresses. The introduction of gradual drought stress is shown to increase rice plants’ propensity to resist against biotic stressors. Genotypes with enhanced ROS-scavenging capabilities such as AC39000 should be prioritised in order to develop rice varieties with dual tolerance to water scarcity and bacterial infection. By selecting these genotypes, it may be possible to develop rice varieties that are more resilient to climate change-induced water stress and the concurrent risk of bacterial blight outbreaks [Garrett et al., 2006], thereby improving food security in vulnerable rice-growing regions. Furthermore, reducing reliance on chemical interventions, such as antibiotics, to control bacterial infections could have significant environmental and economic benefits.

