## Supplementary Data for "Crosstalk Between Drought-Induced ROS Regulation and Resistance to Xanthomonas oryzae Infection in Rice Plants"



October 29, 2024


*

### 1 Shoot Length

#### 1.1 Shoot Length

| Sl. No. | Sample | Shoot Length (cm) |
|---|---|---|
| 1 | HCC1A | 39 |
| 2 | HCC1B | 62 |
| 3 | HCC2 | 54 |
| 4 | HCC3A | 45 |
| 5 | HCC3B | 61 |
| 6 | HCB1A | 54 |
| 7 | HCB1B | 51 |
| 8 | HCB1A | 62 |
| 9 | HCB1B | 44 |
| 10 | HCB3A | 50 |
| 11 | HCB3B | 55 |
| 12 | HGC1A | 39 |
| 13 | HGC1B | 47 |
| 14 | HGC2A | 53 |
| 15 | HGC2B | 57 |
| 16 | HGC3A | 41 |
| 17 | HGC3B | 62 |
| 18 | HRC1A | 36 |
| 19 | HRC1B | 43 |
| 20 | HRC2A | 55 |
| 21 | HRC2B | 49 |
| 22 | HRC3A | 60 |
| 23 | HRC3B | 62 |
| 24 | HCB4A | 46 |
| 25 | HCB4B | 44 |
| 26 | HCB5A | 52 |
| 27 | HCB5B | 47 |
| 28 | HCB6A | 60 |
| 29 | HCB6B | 65 |

| Sl. No. | Sample | Shoot Length (cm) |
|---|---|---|
| 30 | HGB1A | 49 |
| 31 | HGB1B | 63 |
| 32 | HGB1A | 43 |
| 33 | HGB1B | 51 |
| 34 | HGB3A | 57 |
| 35 | HGB3B | 66 |
| 36 | LCC1A | 41 |
| 37 | LCC1B | 48 |
| 38 | LCC2A | 56 |
| 39 | LCC2B | 62 |
| 40 | LCC3A | 43 |
| 41 | LCC3B | 59 |
| 42 | LCB1A | 45 |
| 43 | LCB1B | 62 |
| 44 | LCB1A | 50 |
| 45 | LCB1B | 65 |
| 46 | LCB3A | 57 |
| 47 | LCB3B | 70 |
| 48 | LRC1 | 73 |
| 49 | LRC2 | 42 |
| 50 | LRC3 | 43 |
| 51 | LRB1 | 67 |
| 52 | LRB1 | 40 |
| 53 | LRB3 | 54 |
| 54 | LGB1A | 58 |
| 55 | LGB1B | 61 |
| 56 | LGB1A | 39 |
| 57 | LGB1B | 45 |
| 58 | LGB3A | 62 |
| 59 | LGB3B | 54 |

Table 1: Shoot Length Prior to Bacterial Stress

#### 2 Impact of Bacteria on Chlorophyll Content

| Sample | Day 1 Chl (mg · $g^{-1}$ FW) | Day 5 Chl (mg · $g^{-1}$ FW) | Day 12 Chl (mg · $g^{-1}$ FW) |
|---|---|---|---|
| HCC | 1.687 | 1.684 | 1.724 |
| HCB | 1.454 | 1.306 | 1.166 |
| HGC | 1.704 | 1.569 | 1.553 |
| HGB | 1.469 | 1.307 | 1.212 |
| HRC | 1.591 | 1.544 | 1.556 |
| HRB | 1.475 | 1.354 | 1.259 |
| LCC | 1.255 | 1.249 | 1.235 |
| LCB | 1.249 | 1.105 | 0.966 |
| LGC | 1.371 | 1.343 | 1.337 |
| LGB | 1.161 | 1.045 | 0.959 |
| LRC | 1.166 | 1.161 | 1.165 |
| LRB | 1.169 | 1.024 | 0.944 |

Table 2: Total Chlorophyll Content (Chl) Over Time

#### 3 Impact of Bacteria on ROS Production

| Sample | Day 1 Evans Blue (ng) | Day 1 MDA nmol/mg | Day 5 Evans Blue (ng) | Day 5 MDA nmol/mg | Day 12 Evans Blue (ng) | Day 12 MDA nmol/mg |
|---|---|---|---|---|---|---|
| HCC | 307.0 | 0.006 | 299.0 | 0.006 | 299.0 | 0.007 |
| HCB | 340.0 | 0.041 | 869.0 | 0.051 | 971.0 | 0.061 |
| HGC | 268.0 | 0.007 | 273.0 | 0.009 | 266.0 | 0.008 |
| HGB | 336.5 | 0.041 | 751.5 | 0.054 | 848.5 | 0.067 |
| HRC | 324.0 | 0.008 | 324.0 | 0.008 | 331.5 | 0.007 |
| HRB | 365.0 | 0.052 | 948.0 | 0.057 | 1053.0 | 0.084 |
| LCC | 285.0 | 0.008 | 308.333 | 0.007 | 288.0 | 0.006 |
| LCB | 358.0 | 0.052 | 939.0 | 0.071 | 1030.0 | 0.097 |
| LGC | 287.0 | 0.006 | 292.667 | 0.005 | 294.0 | 0.018 |
| LGB | 349.0 | 0.052419 | 875.667 | 0.065484 | 975.0 | 0.087742 |
| LRC | 368.0 | 0.009435 | 357.5 | 0.033387 | 375.5 | 0.034839 |
| LRB | 355.5 | 0.039194 | 1056.5 | 0.074758 | 1186.5 | 0.101613 |

Table 3: Evans Blue and TBARS Processed Data

### 4 Experimental Setup

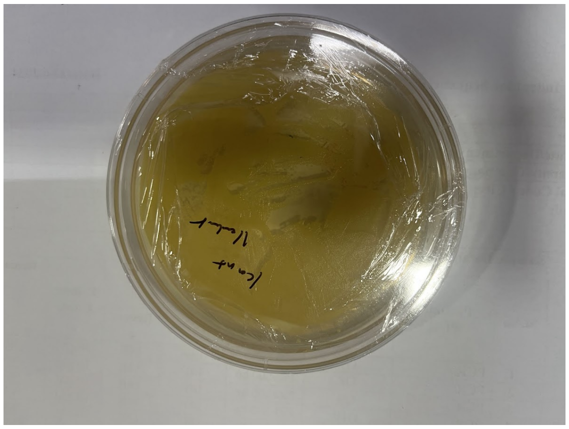

Figure 1: Xanthamonas oryzae on kanamycin-amended agar.

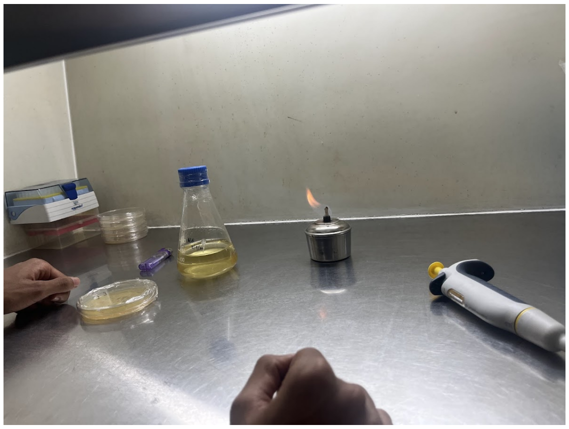

Figure 2: Transfer of bacteria to a nutrient broth shaken at 130 RPM at 32° C.

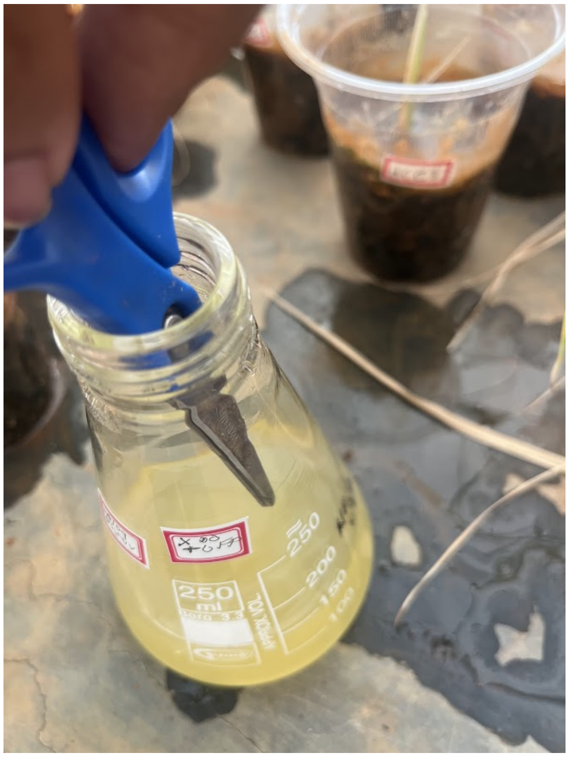


Figure 3: Innoculating the bacteria with the plant by creating an artificial wound.

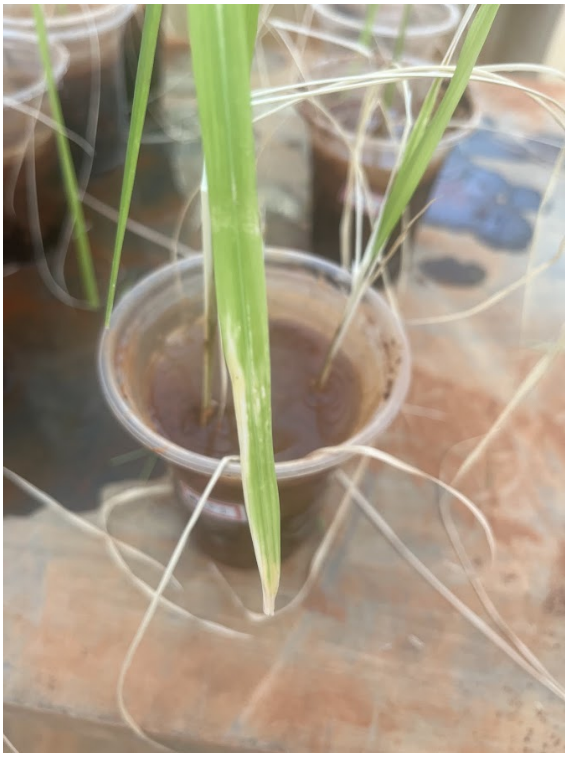

Figure 4: Visible reduction in chlorophyll content due to plant stress.

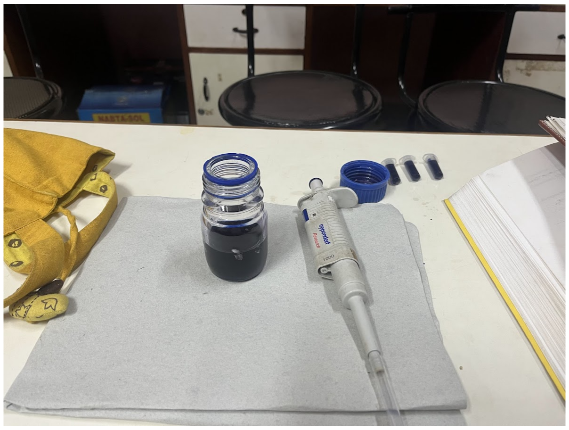

Figure 5: Preparation of 0.25g 100 mL Evans Blue solution.

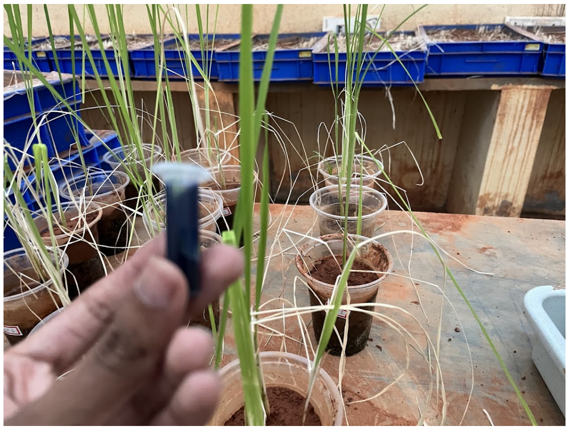

Figure 6: Cut of a leaf sample and storage in eppendorf tube with Evans Blue solution.

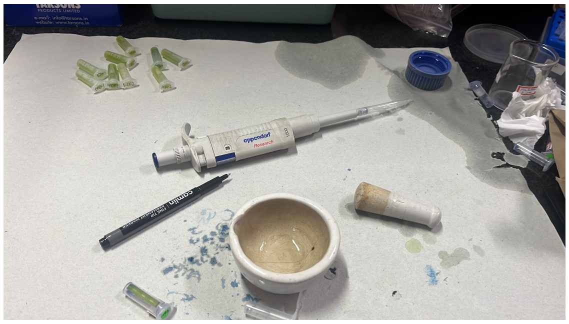


Figure 7: Pestle and mortar to grind and lyse leaf samples in presence of 2 mL of 1% SDS.

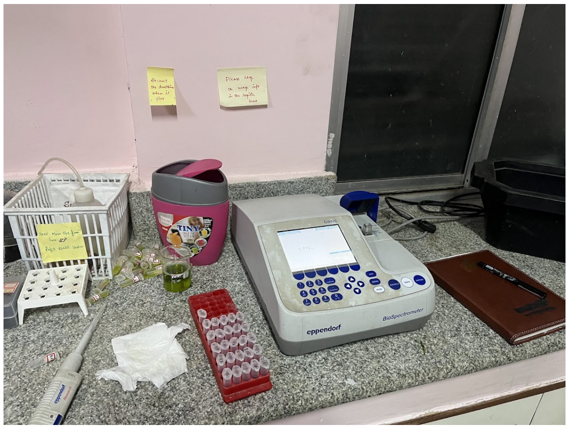


Figure 8: Use of spectrophotometry to assess membrane damage and ROS production.
